# A framework to identify antigen-expanded T Cell Receptor (TCR) clusters within complex repertoires

**DOI:** 10.1101/2021.05.21.445154

**Authors:** Valentina Ceglia, Erin J Kelley, Annalee S Boyle, Yves Levy, Gerard Zurawski, John A Altin

## Abstract

Common approaches for monitoring T cell responses are limited in their multiplexity and sensitivity. In contrast, deep sequencing of the T Cell Receptor (TCR) repertoire offers a global view whose theoretical sensitivity is limited only by the depth of available sampling. However, assignment of antigen specificities within TCR repertoires has become a bottleneck. Here, we combine antigen-driven expansion, deep TCR sequencing and a novel analysis framework to show that homologous ‘Clusters of Expanded TCRs (CETs)’ can be confidently identified without cell isolation, and assigned to antigen against a background of non-specific clones. We show that clonotypes within each CET respond to the same epitope, and that protein antigens stimulate multiple CETs reactive to constituent peptides. Finally, we demonstrate the personalized assignment of antigen-specificity to rare clones within fully-diverse unexpanded repertoires. The method presented here may be used to monitor T cell responses to vaccination and immunotherapy with high fidelity.

## Introduction

The identification, within complex repertoires, of T cells specific for a target of interest is an essential immunological capability, used to diagnose infection^1^ (Nyendak et al., 2009) and measure the immunogenicity of vaccines and immunotherapies^2^ (Flaxman and Ewer, 2018). Current methods for quantifying rare antigen-specific T cells include assays that measure antigen-stimulated cytokine production (e.g., immunospot assays and flow cytometric detection of intracellular cytokines^3,4^ (Lovelace and Maecker, 2011; Sidney et al., 2020)), as well as assays that use labeled peptide:MHC probes to directly detect antigen-binding T cells^5^ (Altman et al., 1996). Although widely useful, the ability to multiplex these assays across targets is limited, as is their sensitivity to detect rare T cell responses.

T cells recognize MHC-restricted peptide antigens by means of the heterodimeric T Cell Receptor (TCR), encoded by somatically-diversified □ and β loci^6^ (Davis et al., 1984). The rearranged TCR α:β sequence pair completely determines a T cell’s specificity, and current technologies enable >10^7^ unpaired or >10^4^ paired TCR chains to be routinely sequenced from a sample^7^ (Yost et al., 2020). In contrast to traditional methods of antigen-specific T cell detection, deep sequencing of TCRs can reveal complete repertoires with high sensitivity. However, the ability to confidently assign antigen reactivities to (or ‘decode’) particular TCR sequences within this repertoire has become a bottleneck.

One approach to decoding the repertoire, ‘exposure association’, involves associating the incidence of particular clonotypes (e.g., defined at the CDR3β amino acid sequence level) with antigen exposure status within a cohort of individuals. This approach has the potential to reveal ‘public’ sequences that are enriched in exposed subjects, and has been used to accurately classify cytomegalovirus (CMV) serostatus^8^ (Emerson et al., 2017), and more recently to diagnose SARS-CoV-2 infection^9^ (Snyder et al., 2020). The ability to discover antigen-associated public clonotypes has powerful diagnostic potential, however discovered associations have generally been too weak to allow high-confidence assignment of antigen-specificity to particular public clonotypes within any given individual. This approach is also limited by a requirement for large cohorts of exposed and unexposed individuals to identify sequences with statistical confidence.

A second approach, ‘probe association’, involves the use of probes to isolate T cells that recognize defined antigens within particular samples. Multimerized peptide:MHC probes have been used for decades to identify and isolate T cells in an antigen-resolved fashion^5^ (Altman et al., 1996). Methods combining antigen restimulation with the detection of upregulated cellular response markers can also been used for this purpose^10,11^ (Klinger et al., 2015; Dan et al., 2016). Although these approaches allow a powerful interrogation of the T cell response, the rarity of antigen-specific cells against non-specific background binding renders some memory T cell responses below the limit of detection, and the peptide:MHC multimer approach depends on the *a prior* identification of appropriate peptide:MHC combinations.

Thirdly, ‘sequence-based prediction’ describes a new family of methods in which growing catalogs of defined TCR:antigen combinations are used to train machine learning algorithms to predict specificity directly from TCR sequences^12,13,14^ (Bagaev et al., 2020; Fischer et al., 2020; Tong et al., 2020). These have great potential to enable generalizable decoding of the repertoire, especially as the training datasets grow, however they do not yet enable the confident assignment of specificities within deep repertoires using TCR sequences alone.

Here, we develop an alternative approach to decoding TCR repertoires. In this method, rare T cells are clonally expanded by antigens of interest in culture, subjected to bulk TCR sequencing, and clonal frequencies are analyzed using a similarity-based clustering approach to identify and organize families of antigen-responsive clonotypes against the majority of irrelevant sequences.

## Results

### Optimizing the conditions for whole protein antigen-driven *in vitro* expansion of T cells

The model we used for this study was the steady state memory T cell repertoire specific to Influenza matrix protein (M1) in healthy adult donors. M1 is >90% conserved across strains and dominates the cross-reactive memory CD4^+^ and CD8^+^ T cell repertoire in healthy individuals^15^ (Lee at al., 2008). Thus, repeated seasonal infections and vaccination account for the presence of M1-specific T cell memory in most healthy donors^16,17^ (Terajima et al., 2008; Furuya-Kanamori et al., 2016). We and others have demonstrated that linking peptide or whole protein antigens to antibodies directed to Dendritic Cell (DC) receptors such as CD40 can efficiently potentiate antigen-presentation, resulting in efficient expansion of both CD4^+^ and CD8^+^ T cells across multiple epitopes and HLA specificities within *in vitro* culture systems^18,19,20^ (Bozzacco et al., 2007; Flamar et al., 2012, 2013). We have previously described a convenient method for non-covalent assembly of anti-Dendritic Cell (DC) antibodies and antigens using a bacterial dockerin (doc) domain fused to the antibody heavy chain C-terminus, and antigen such as Flu M1 fused to a cohesin (coh) counter-domain^19^ (Flamar et al., 2012).

We cultured Peripheral Blood Mononuclear Cells (PBMCs) obtained by apheresis of normal donors (ND) with dose ranges of a cohesin-Flu M1 fusion protein alone (Flu M1) or in complex with three different CD40-targeting antibody vehicles. After an expansion culture period of 10 days, cells were harvested and re-stimulated with 3 pools of overlapping 15 mer peptides covering the entire Flu M1 protein, then analyzed by Intracellular Cytokine Staining (ICS) for peptide-elicited production of intracellular IFNγ and TNFα. Figure 1 shows that in ND1004, Flu M1-specific CD4^+^ T cells from epitopes within all three M1 regions were elicited with the CD40-targeted antigen being 10-100-fold more efficacious than Flu M1-stimulation alone. Up to 20% of the T cells in the 10-day culture with 0.1 nM anti-CD40 11B6-CD40L:Flu M1 stimulation produced IFNγ and/or TNFα specifically in response to Flu M1 peptides versus <1% elicited by 0.1 nM untargeted Flu M1. This is consistent with other data showing the high *in vitro* efficiency of targeting antigens to CD40 in PBMC cultures^18,21^ (Flamar et al., 2013; Yin et al., 2016). In contrast, in this donor Flu M1-specific CD8^+^ T cells were not significantly expanded in any of the conditions (Figure 1a) with responses below 2% and no clear trends related to stimulation condition or dose.

**Figure 1.**
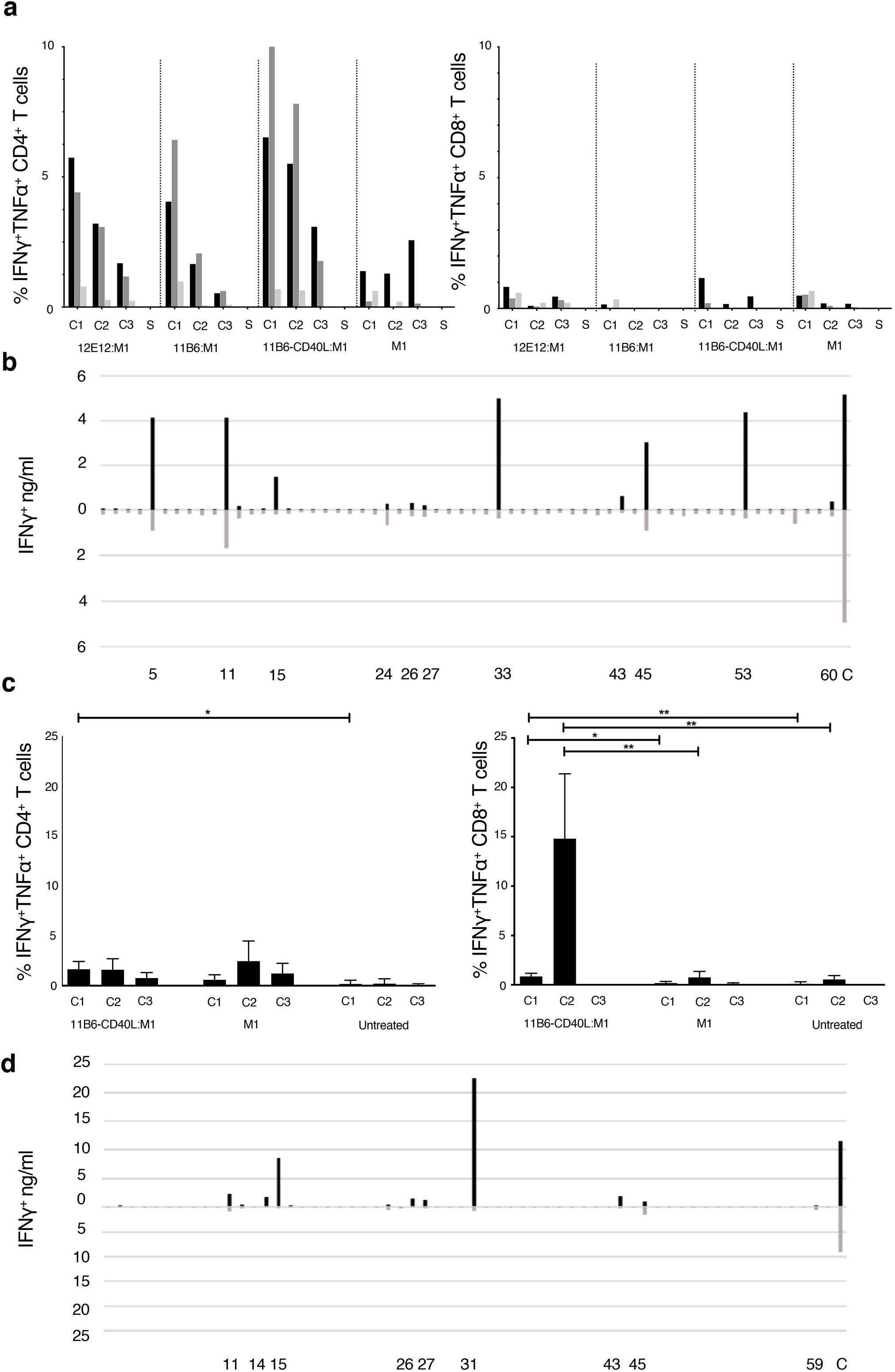
(a, c) Expansion of epitope-reactive T cells by M1 protein formulations. Analysis of Flu M1-specific T cell responses in a day 10 ND1004 (a) and ND1005 (c) PBMC culture stimulated with CD40-targeted Flu M1 protein. The left panel shows the % of IFNγ^+^ and/or TNFα^+^ antigen-specific CD4^+^ T cells as determined by ICS analysis. Baseline values for solvent (S, no peptide) controls were subtracted. In (a) cultures contained a dose range of CD40-targeting antibodies conjugated to cohesin Flu M1 or Flu M1 alone (M1), while in (c) only one dose was tested. The right panel shows analogous data for % of IFNγ^+^ and/or TNFα^+^ CD8^+^ T cells. In (a) baseline S values for the CD4^+^ were 0.43±0.14%, and for the CD8^+^ were 0.8±0.4%. Compared to a starting input of PBMCs, the end stage 10 day cultures increased in total numbers as follows for, respectively, the 1, 0.1, and 0.01 nM conditions: anti-CD40 12E12:M1 5.6, 3.6,1.6-fold; anti-CD40 11B6:M1 12.5, 2.5, 0.9-fold; anti-CD40 11B6-CD40L:M1 7, 2.7 1-fold; and M1 1.5, 0.9, 0.8-fold. In (c) the data show results from 4 independent experiments with ND1005. Values for solvent without peptide stimulation (S) were subtracted from each peptide stimulation point; baseline S values for the CD4^+^ between 0.1 and 1, and for the CD8^+^ between 0.3 and 2. Cells after 10 day expanded 6.6±2.4 fold with anti-CD40 11B6-CD40L:M1 and 3.3±1.2 fold with M1 alone compared to cells alone. * P ≤ 0.05, ** P ≤ 0.01. The CD4^+^ T cell two-tailed T test comparison is between data for anti-CD40 11B6-CD40L:M1 and cells alone; there were no other significant differences in the CD4^+^ T cell responses. The CD8^+^ T cell two-tailed T test comparison is between data for anti-CD40 11B6-CD40L:M1 and Flu M1; there were no significant differences in the responses to Flu M1 compared to cells alone. **(b, d)** Flu M1 peptide-specific T cell repertoire of ND1004 (b) and ND1005 (d) expanded by anti-CD40-11B6-CD40L Flu M1 targeting.

To ascertain the breadth of the expanded Flu M1-specific T cell responses elicited by targeting Flu M1 with anti-hCD40 11B6-CD40L, day 10 cultures were re-stimulated with individual 15 mer Flu M1 peptides and IFNγ secretion was measured 48 hours later. Figure 1b shows that at least 10 Flu M1 peptide specificities were elicited by anti-CD40 11B6-CD40L:Flu M1 targeting and many of these were also detected at lower response levels by non-targeted cohesin-Flu M1.

PBMCs from a second normal donor (ND1005) were cultured with 1 nM anti-CD40 11B6-CD40L:Flu M1 complex or 1 nM cohesin-Flu M1 alone, and after an expansion culture period of 10 days, cells were harvested and re-stimulated with 3 pools of overlapping 15 mer peptides covering the entire Flu M1 protein, then analyzed by ICS for peptide-elicited production of intracellular IFNγ and TNFα. Figure 1c shows that in this donor anti-CD40 11B6-CD40L:Flu M1 complex elicited a low level but significant ∼1% M1-specific CD4^+^ T cell response from epitopes within the C1 Flu M1 region. However, in replicate experiments, 8-21% of the CD8^+^ T cells in culture with 1 nM 11B6-CD40L:Flu M1 stimulation produced IFNγ and/or TNFα specifically in response to Flu M1 C2 peptides versus <1.5% elicited by 1 nM untargeted cohesin-Flu M1. The breadth of the expanded Flu M1-specific T cell responses elicited by anti-CD40 11B6-CD40L:Flu M1 and Flu M1 alone were determined in day 10 cultures re-stimulated with individual 15 mer Flu M1 peptides and then assayed for IFNγ secretion after 48 hours. Figure 1d shows that at least 8 Flu M1 peptide specificities were elicited by anti-CD40 11B6-CD40L:Flu M1 targeting and most of these were also detected at generally lower response levels by untargeted cohesin-Flu M1.

### Identification of antigen-expanded clonotypes within the repertoire

The above experiments established that the two selected donors contain a broad repertoire of memory Flu M1-specific CD4^+^ T cells (ND1004) and CD8^+^ T cells (ND1005) that could be efficiently expanded *in vitro* from 10 day PBMC cultures stimulated with low doses of Flu M1 targeted to CD40 on APCs, especially via the anti-CD40 11B6-CD40L antibody vehicle.

To profile the TCR repertoire, we extracted RNA from the cultured cells of these donors, generated a library of full-length RNA products, performed nested PCR enrichment by priming against the *TRA* and *TRB* constant regions, and sequenced the resulting amplicons. Across the 5 conditions (no-antigen, Cohesin-Flu M1 protein at 0.1 nM or 1 nM, or anti-CD40 11B6-CD40L:Flu M1 at 0.1 nM or 1 nM), we recovered a total of 187,250 and 50,762 TCRα and 124,120 and 43,607 TCRβ productively-rearranged clonotypes (each defined as a unique combination of the CDR3 nucleotide sequence and mapped V+J segments) from ND1004 and ND1005, respectively. Strikingly, 86-94% of all detected TCRα and TCRβ clonotypes were unique to a particular culture. Moreover, even when comparing the no-antigen condition (hereafter ‘Ag–’) against the most stimulatory condition (anti-CD40 11B6-CD40L:Flu M1 at 1 nM, hereafter ‘Ag+’), there was no consistent evidence of a strong antigen-driven effect on the overall clonal frequency distributions. For example, of all clonotypes detected in either the Ag– or Ag+ condition for ND1004, 39% and 57% were uniquely present in the respective condition and absent in the other. For ND1005, these numbers were 15% and 81%. Together, these observations indicate strong culture-specific effects on clonal frequencies that are independent of the added antigen, precluding the confident assignment of antigen-specificity to clonotypes based on an analysis of frequencies alone.

To increase the power to identify antigen-expanded clonotypes, we reasoned that stimulation with antigen should expand families of clones that use homologous TCRs to recognize the same peptide:MHCs^22,23^ (Dash et al., 2017; Glanville et al., 2017). However, unlike T cells purified according to reactivity to individual antigens, we expect clonal families in expanded cultures to be admixed within a majority of irrelevant clones. To identify such families, we developed a method (Figure 2) based on clustering of the 1000 most frequent TCR sequences (αs and βs separately) in a sample using comprehensive pairwise homology measurements.

**Figure 2.**
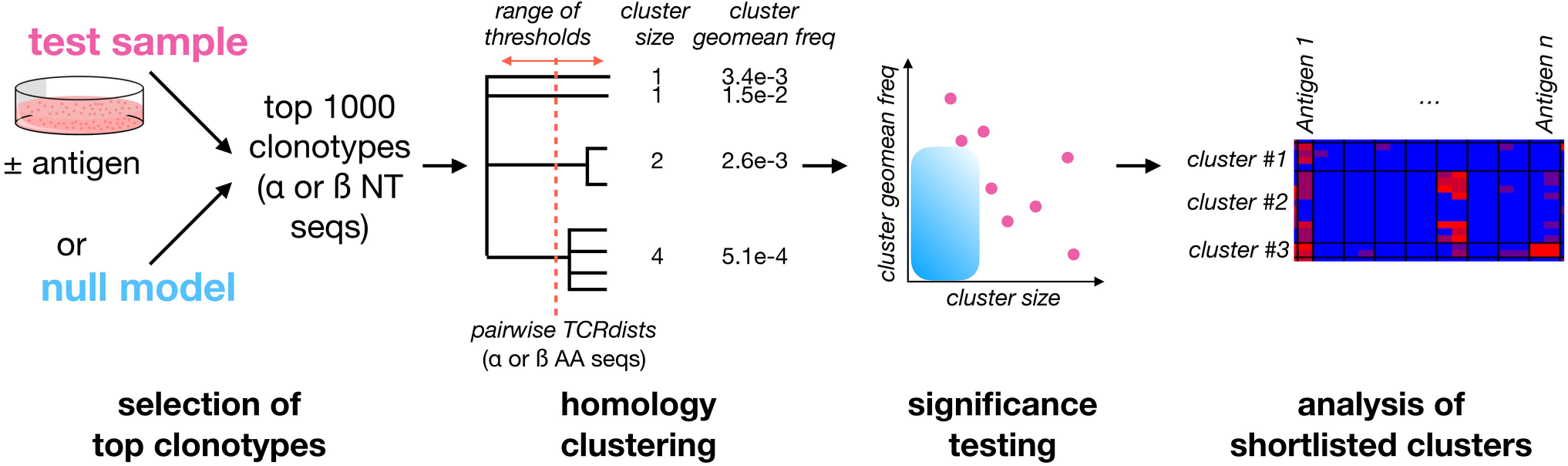
A framework for identifying Clusters of Expanded TCRs (CETs) within complex repertoires. To identify antigen-responsive clonotypes admixed within a large population of irrelevant clonotypes, the 1000 most abundant TCRα or TCRβ clonotypes for the sample of interest (resolved at the nucleotide level and quantified by deep sequencing) are analyzed by similarity-based clustering of their CDR amino acid sequences. Comprehensive pairwise similarity measurements using the TCRdist metric are used to identify clonotype clusters across a range of thresholds. The significance of each Cluster of Expanded TCRs (CET) is then quantified as the probability of observing a cluster with the same number of members, at or above its observed mean frequency, within trials of 1000 randomly-selected clonotypes. Finally, shortlisted clonotypes are analyzed for their abundance across multiple conditions.

For each sample we implemented the recently-described TCRdist metric^22^ (Dash et al., 2017) – which provides a quantitative measure of amino acid similarity between the exposed CDR loops of any 2 TCRs – across all possible pairs among the 1000 most frequent TCRα or TCRβ clonotypes, resulting in ∼1e6 total comparisons. Clusters of expanded TCRs (‘CETs’) were then identified at a range of similarity thresholds, and each CET parametrized according to (i) its number of members and (ii) their geometric mean frequency within the overall TCR population. To exclude CETs that could occur by chance, we next estimated the significance of each CET by determining how commonly a cluster with the same number of members, and equal or greater mean frequency, arises at the same threshold based on a set of 1000 randomly-generated TCR sequences drawn from a matched underlying frequency distribution. The sliding threshold approach is designed to enable sensitivity to TCR groups across the size:frequency spectrum: ranging from high-frequency TCR groups with few members, to lower frequency groups with more members. While insensitive to antigen-specific clonotypes that do not form homology clusters, this method uses the statistical power of convergent antigen recognition to allow antigen-expanded TCRs to be confidently identified within individual samples (without an intrinsic dependency on controls or replicates), enabling condition-specific hypotheses to be tested subsequently with greater power on a more focused set of clonotypes.

Using TCRdist thresholds ranging from 0-50, we applied our clustering method to the TCRα and TCRβ clonotypes sequenced in the Ag– versus Ag+ conditions for ND005, as well as to a randomly-generated control set of clonotypes. The total number of detected CETs was greater in the Ag+ compared to Ag– condition, and was lowest in the randomly-generated set (Figure 3b, left). When focusing only on CETs that passed the significance test (described above, based on cluster sizes and mean frequencies relative to a random model), the enrichment in the Ag+ condition was more marked, and as expected, none of the clusters detected in the random control reached significance for any TCRdist threshold (Figure 3b, right). CET analysis therefore revealed an asymmetry between the Ag– versus Ag+ conditions that is expected, yet much less obvious on a frequency-only analysis.

**Figure 3.**
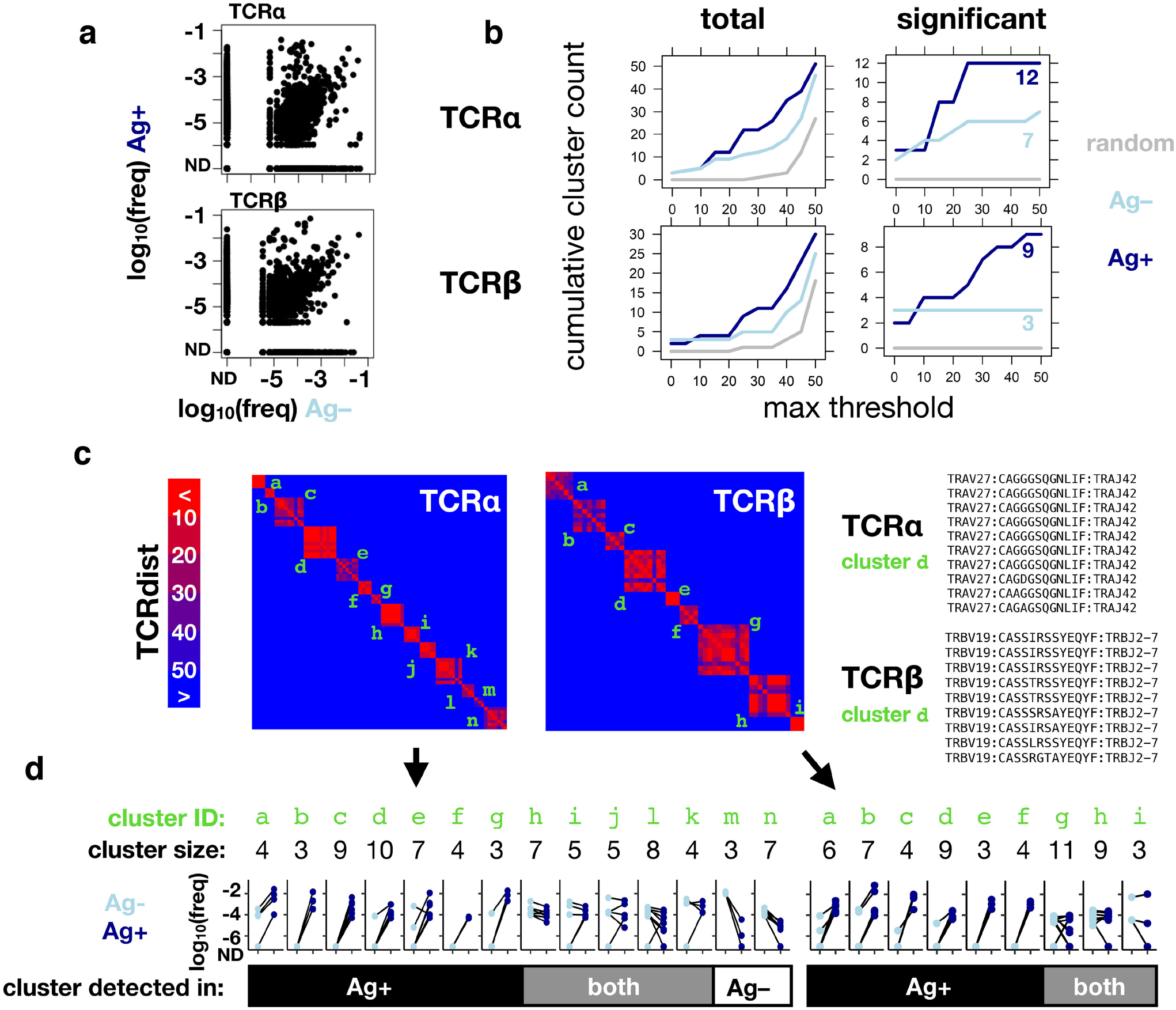
CET analysis enables identification of antigen-responsive clonotypes within complex repertoires. PBMCs cultured with IL-2 for 10 days, in the presence (Ag*+*) or absence (Ag*–*) of influenza M1 protein formulations (as described in Figure 1), after which RNA was extracted for amplification of TCRα and TCRβ chains, sequenced and analyzed using the scheme shown in Figure 2. This figure shows data for one representative donor (ND1005) and antigen (1 nM of the anti-CD40 11B6-CD40L:Flu M1). (**a**) Frequencies in Ag+ and Ag– conditions for all detected clonotypes. (**b**) Cumulative number of CETs – either total (*left*) or significant (*right*) – detected up to the threshold shown on the x-axis. For comparison across thresholds, clonotypes within CETs detected up to each threshold were combined and re-clustered at the maximum threshold prior to enumeration. (**c**) Map showing pairwise TCRdist similarities of clonotypes within significant TCRα and TCRβ CETs from the analysis described in **b**. Identifiers in green indicate CETs detected up to the maximum threshold of 50. V segment, CDR3 and J segment sequences are shown for detected clusters that corresponding to a well-known HLA-A2-restricted M1 reactivity (*right*). (**d**) Clonotype frequencies in the Ag*+* and Ag*–* cultures for members of each of the significant TCRα and TCRβ CETs.

Since our method looks only for TCR clusters expanded within an individual sample, without regard to the presence or absence of antigen, the CETs identified in the Ag+ condition could represent clonotypes expanded either *in vivo* (against any antigen) or *in vitro* (against Flu M1). We resolved between these possibilities by comparing the identities and frequencies of the significant CETs detected between the Ag+ and Ag– conditions. Combined across both the Ag+ and Ag– conditions, the analysis revealed 14 significant TCRα CETs, comprising 3-10 members (79 total clonotypes), and 9 significant TCRβ CETs, comprising 3-11 members (56 total clonotypes) (Figures 3c,d).

Strikingly, 5/7 and 3/3, respectively, of the α and β CETs detected in the Ag– condition were also significant in the Ag+ condition, and their constituent clonotypes generally showed minimal differences in frequency between the 2 conditions, indicating their expansion independently of the Flu M1 antigen and likely *in vivo* prior to culture. In contrast, the majority of CETs (7/12 and 6/9 α and β, respectively) detected in the Ag+ condition were not detected in the Ag– condition, reflecting the fact that their constituent clonotypes were dramatically (100-10,000-fold) expanded in the Ag+ condition.

The largest TCRα CET contained 10 clonotypes, each comprising the *TRAV27*/*TRAJ42* segment pair with consensus CDR3 sequence ‘CAGxGSQGNLIF’. Similarly, the largest TCRβ cluster detected only in the Ag+ condition contained 9 clonotypes, each comprising the *TRBV19*/*TRBJ2-7* pair and with CDR3 consensus CASSxRSSYEQYF (Figure 3c, right). These correspond precisely with a known TCRa:β paired motif previously described for CD8^+^ T cells recognizing the immunodominant HLA-A2-restricted Flu M1 peptide GILGFVFTL^22^ (Dash et al., 2017), consistent with ND005’s status as HLA-A2+ and validating the algorithm’s ability to robustly identify an expected antigen-specific TCR clonotype within the unpurified repertoire.

### Characterization of TCR clusters across distinct antigens and formulations

To test the hypothesis that the detected CETs correspond to groups of TCRs united by their antigen recognition at the epitope-level, we reasoned that the members of each CET should respond in a co-ordinated way when expanded with different constituent antigenic peptides from the protein antigen. We identified candidate Flu M1 antigenic peptides in our study subjects from the patterns of re-stimulated cytokine production shown in Figure 1b and d, and used these to generate additional cultures in which T cells from the same donors were stimulated with 2 μM of either single or small clusters of overlapping Flu M1 peptides or polyclonal stimulation with phytohemagglutinin (PHA) as positive control. After 10 days of culture, the cells were re-stimulated with the matching peptides for 48 hours or with PHA (C) and the collected supernatants were analyzed for IFNγ production (Figure 4). RNA from these cultures, together with the original ones in which T cells from the same donors were cultured with the CD40-targeted or untargeted whole Flu M1 protein, was then analyzed for the representation of TCRα and TCRβ clonotypes.

**Figure 4.**
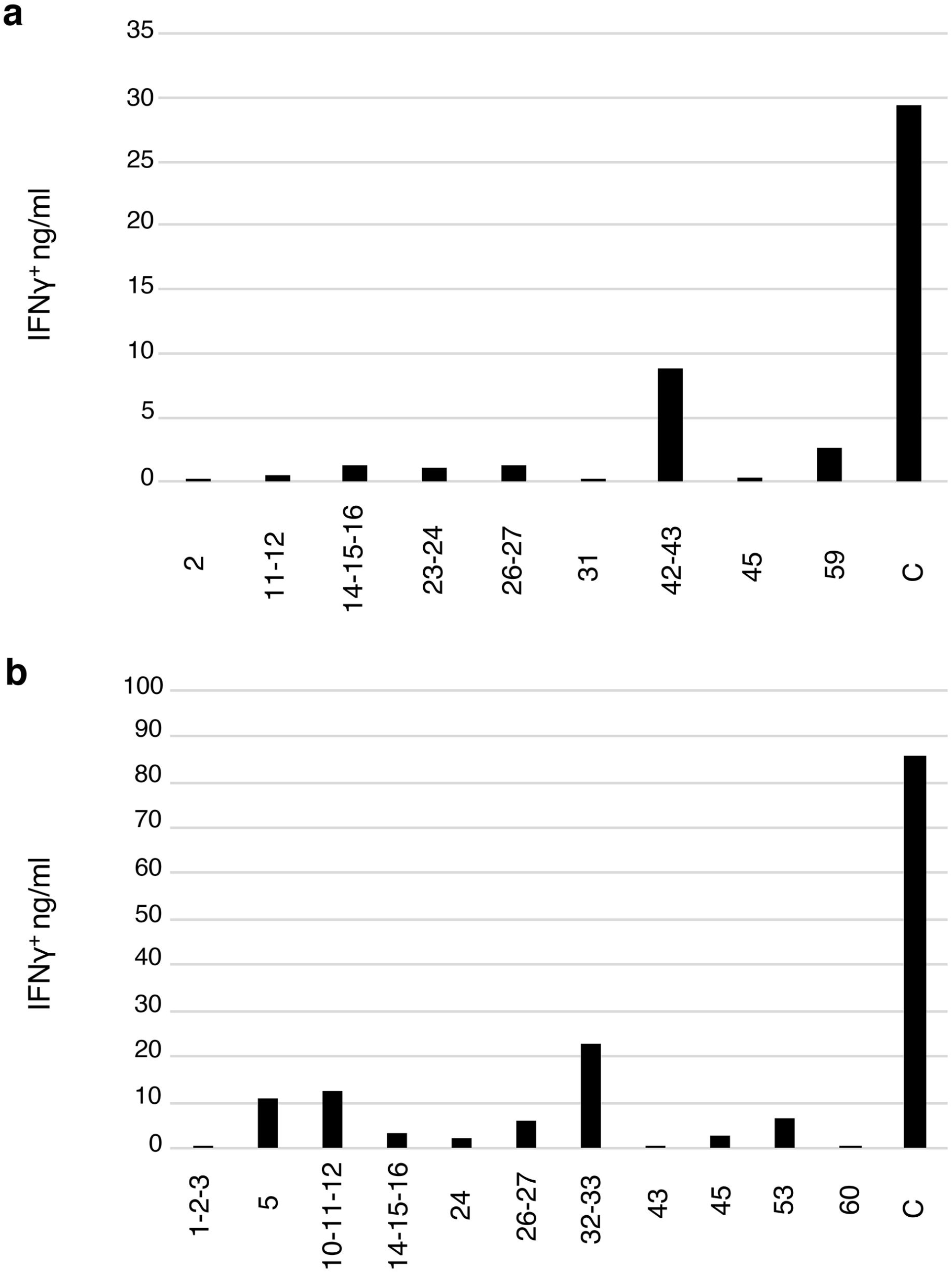
Flu M1 epitope-specific T cell repertoires expanded by single or small clusters of Flu M1 peptides. **(a)** ND1005 and (**b**) ND1004 PBMCs were cultured in complete RPMI 1640 + 10% AB serum for 10 days in the presence of IL-2, stimulated with 1 μM of either single peptides or small clusters of Flu M1 peptides. At day 11 cultures were re-stimulated with 2 μM the matching Flu M1 peptides or PHA as a control for 48 h and the collected supernatants were analyzed for IFNγ.

We applied the CET-detection algorithm described above to TCRα and TCRβ clonotypes from each condition in the 2 donors, and aggregated the identified clonotype clusters across the different conditions to generate a master list for each donor that was then re-clustered for display. TCRα and β CETs whose members show an average expansion of at least 100-fold in any condition over the Ag– condition are shown in Figure 5. In total, we identified 11 and 8 antigen-enriched TCRα and TCRβ CETs (comprising, respectively, 88 and 70 total clonotypes) meeting those criteria in ND004, and 31 and 16 TCRα and TCRβ CETs (comprising, respectively, 207 and 84 total clonotypes) in ND005.

**Figure 5.**
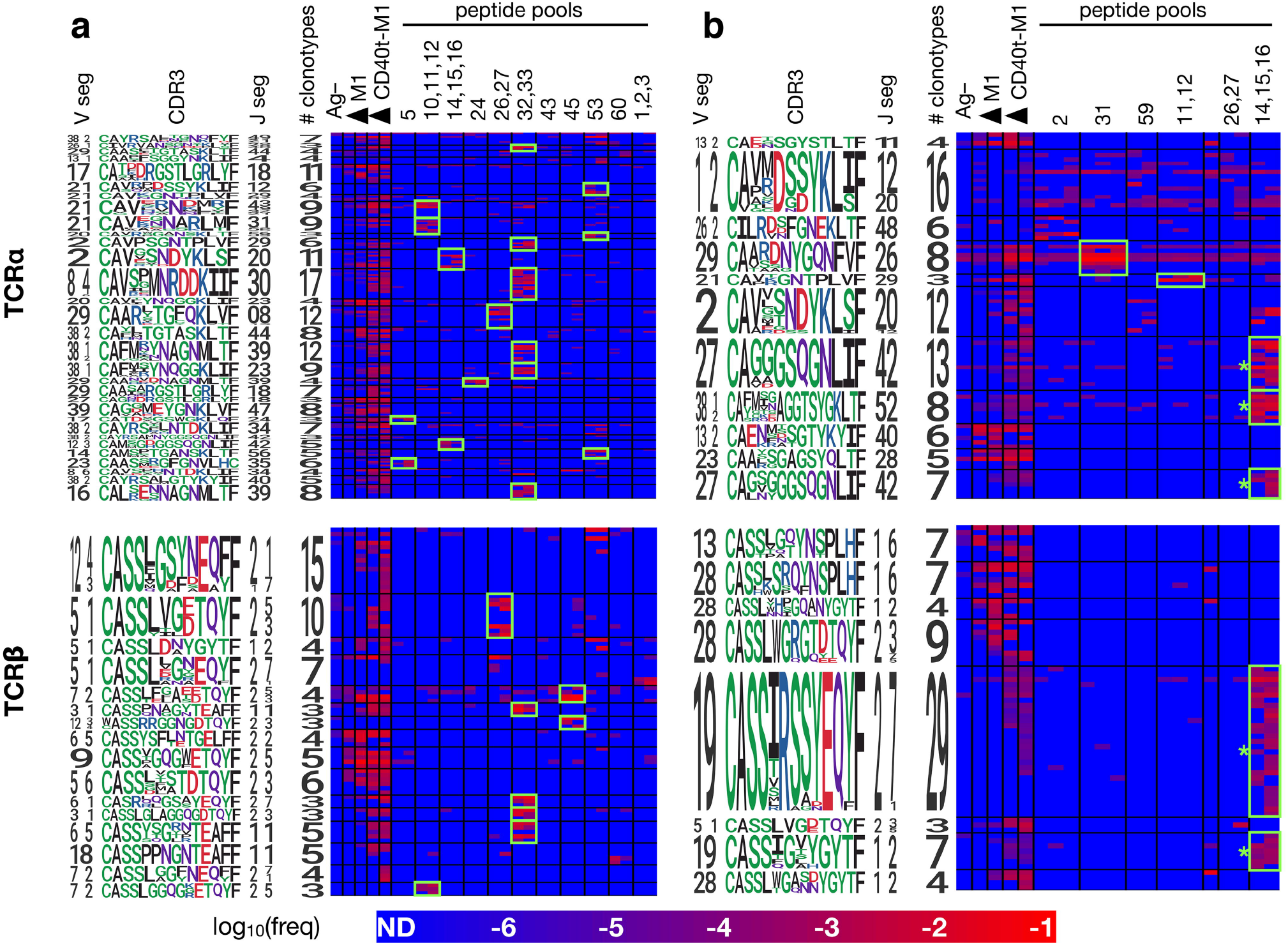
Members of each CET show co-ordinated responses that distinguish different forms of antigen. PBMCs from ND004 and ND005 (**a** and **b**, respectively) were expanded in replicate cultures with influenza M1 antigen in a variety of forms – untargeted or CD40-targeted whole protein, or pools of overlapping peptides corresponding to the reactive epitopes identified in Figure 4, and then analyzed by TCR sequencing and CET identification as described previously. Significant CETs were identified in each sample individually, aggregated across all samples, and then re-clustered at the maximum TCRdist threshold of 50 for display. Each row represents a single clonotype, with CETs demarcated by horizontal black lines and labeled by logos representing their constituent V, CDR3 and J sequences. Each column represents a single culture, with conditions demarcated by vertical black lines (one replicate per column). Shown are CETs with ≥3 members and ≥100X average enrichment in any condition over the Ag– condition; highlighted in green are CET:peptide combinations with ≥100X average enrichment over the Ag– condition. * = CETs whose sequence features closely match TCRs previously described to recognize the HLA-A2-restricted GILGFVFTL antigen^22^ (Dash et al., 2017).

**Figure 6.**
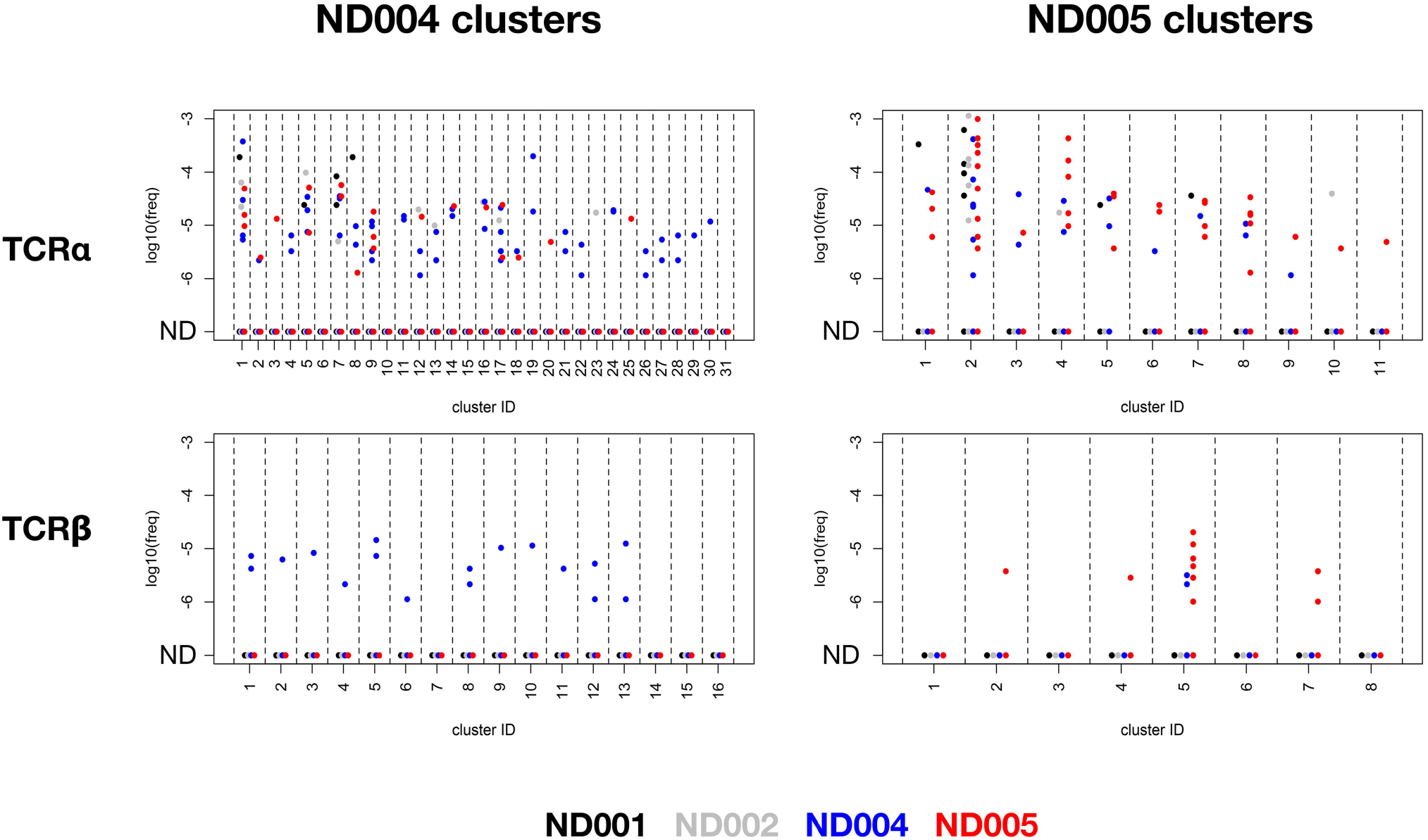
Specificity of rare antigen-specific TCRβ, but not TCRα, clonotypes within deep unenriched repertoires. TCR α and β libraries were prepared from uncultured PBMCs from 4 healthy subjects (ND001, ND002, ND004 and ND005), and deeply sequenced to generate an average of >1.3M clonotype counts per sample. Clonotypes within each of the TCRα and β CETs identified previously in ND004 and ND005 (Figure 5) were queried against these 4 deep, unexpanded datasets by matching for nucleotide-level sequence identity. Plots show log10(frequency) of sequences in the 4 unexpanded datasets (colored by donor) and organized by chain type (upper / lower), CET donor (left / right) and CET grouping (x-axis groups).

Consistent with our hypothesis, clonotypes members within the identified clusters showed patterns of reactivities across the different Flu M1 antigens that were strongly-co-ordinated. In both donors, the anti-CD40 11B6-CD40L:Flu M1 formulation expanded the largest number of CETs, consistent with its enhanced immunogenicity compared to untargeted protein, and reflecting the cytokine production patterns that we observed (Figure 1). For each of the 4 donor:TCR chain combinations, the untargeted cohesin-Flu M1 protein expanded a significantly smaller group of CETs, in each case being a subset of those expanded by the targeted version of the protein. Consistent with expectations, a majority (11/17) of the peptide pools expanded at least 1 (and up to 6) discrete CETs, and there was a general correlation between the number of α v β CETs across donor/antigen combinations, the most striking example being peptide pool 32,33 which expanded 6 α and 4 β CETs in ND004. Segment usage and CDR3 motifs were largely non-conserved across these CETs, suggesting that the peptide pool contains multiple (but nearby) epitopes, and/or that TCRs with diverse sequence features are recognizing the same peptide:MHC complex.

Conversely, cluster specificity at the peptide level was evident from the fact that a CET never responded to more than 1 distinct peptide pool. Moreover, the ‘14,15,16’ peptide pool, which covers the immunodominant HLA-A2-restricted epitope ‘GILGFVFTL’ mentioned previously, stimulated 5 CETs (3 α and 2 β – denoted by ‘*’ in Figure 5b) all of which correspond to previously-described TCR sequence motifs for this epitope^22^ (Dash et al., 2017). Interestingly, these CETs contained an unusually large number of clonotypes (28 TCRα and 36 TCRβ) and were most strongly expanded in the peptide-only condition, followed by the targeted-protein condition, and not significantly expanded at all in the untargeted-protein condition. This observation suggests limitations in antigen processing in the case of the whole-protein antigen, and that these might be overcome by the CD40-targeting.

### Quantification of antigen-specific clonotypes within matched unexpanded repertoires

A key motivation for developing methods for decoding TCR repertoires is to enable multiplexed and sensitive monitoring of rare T cells. Having identified sets of high-confidence Flu M1-responsive TCR clonotypes from study subjects ND004 and ND005, we next tested whether these responses could be detected in their unexpanded states in deep *ex vivo* samples.

Since it is theoretically possible that an unpaired TCR chain detected in an individual in fact derives from multiple TCRs of different specificities (owing to pairing with different chain partners), we measured the frequency with which clonotypes identified by the CET analysis are observed within deep unexpanded samples from both matched and unmatched donors. We reasoned that detection of these TCR clonotypes in samples from the matched, but not the unmatched, donors would indicate assay specificity, and set a limit on the frequency with which chain rearrangements could converge by chance and confound the analysis. Conversely, occurrence of the queried sequence in unmatched donors would indicate a specificity limit, beyond which the inferred link between unpaired clonotype sequence and antigen specificity may break down.

To that end, we sequenced uncultured PBMCs from a total of 4 healthy donors – the 2 characterized so far (ND004, ND005), and 2 additional controls (ND001 and ND002) – to an average depth of 1.3e6 mapped TCR clonotype reads. We then queried these 4 deep unexpanded repertoires for nucleotide-level sequence matches across each of the 295 α and 154 β clonotypes contained in the Flu M1-responsive CETs that we previously identified in ND004 and ND005 (using the analyses described in Figure 5).

For the Flu M1-responsive TCRαs, ≥1 clonotype was detectable in ≥1 of the 4 unexpanded samples in a large fraction of CETs (27/31 and 11/11 of the ND004 and ND005 CETs, respectively), with frequencies ranging from 1e-6 to 3e-3. While these detectable unexpanded clonotypes occurred disproportionately in the samples matching the donors from which they were identified (48/83 and 35/67 for the ND004 and ND005 clonotypes, respectively), there was also a substantial fraction of occurrences in unmatched donors, across a similar range of frequencies.

For TCRβs, in contrast, the overall matching rates (≥1 clonotype was detectable in ≥1 donor in 12/16 and 4/8 ND004 and ND005 CETs, respectively) and frequencies (1e-6 to 5e-4) appeared somewhat reduced, but now these clonotypes were highly-specific for the matching donor. Among the Flu M1-responsive TCRβ ND004 clonotypes that were also identified in an unexpanded sample, all 17/17 were identified in ND004. For ND005, this rate of ‘matching hits’ was 10/12, with 2 ‘unmatched’ clonotypes in cluster #5 (corresponding to TCRs recognizing the HLA-A2-restricted ‘GILGFVFTL’ epitope) detected in ND004. The antigen-specificity of these 2 clonotypes is unknown, and it remains possible that they also recognize this same immunodominant epitope. Overall, we conclude that TCRβ, but not TCRα, clonotypes assigned to antigen using the CET analysis are often detectable with high specificity in the unexpanded state, down to frequencies ∼1e-6.

## Discussion

The approach described here allows TCR sequences within complex repertoires to be confidently assigned to antigens of interest. Unlike existing approaches that require cell labeling and isolation, our method uses a statistical analysis of deeply-sequenced TCR repertoires in response to antigen-driven expansion. Taking advantage of the fact that individual epitope-specific immune responses often comprise groups of homologous TCRs, we integrate both TCR frequency and sequence homology information across the repertoire to identify groups of antigen-expanded clonotypes within individual samples. Using this method, we observe that the response to a model protein antigen comprises groups of homologous clones raised against distinct peptide epitopes, and that many of the responding TCRs are also detectable in their rare, unexpanded state in *ex vivo* samples. We also show that the number of antigen-specific clonotypes detected can be dramatically augmented by CD40-targeting.

Consistent with previous work, our analysis demonstrates that the peptide:HLA-specific T cell response within a donor frequently comprises a large number of individual clonotypes whose TCRs use convergent sequence features to recognize the antigen. The most striking example observed here is the 29-member *TRBV19+*/*TRBJ2-7+* cluster recognizing the well-documented immunodominant HLA-A*02:01-restricted GILGFVFTL peptide. The factors contributing to the activation of such a large T cell family within an individual also likely underlie immunodominance *across* individuals – namely, a high generation probability of T cell precursors capable of recognizing the antigen, and abundant or sustained expression of the antigen during infection^4,24^ (Sidney et al., 2020; Oseroff et al., 2008).

The expansion-based approach described here differs in several notable ways from alternative methods for TCR mapping that use cell labeling and isolation. At a technical level, the method herein does not rely on cell isolation, nor does it require peptide:MHC multimer probes to be identified and constructed. It does, however, involve antigen expansion cultures, which may become a bottleneck when interrogating large numbers of antigens. However, like existing approaches, it is likely that the number of targets analyzed simultaneously can be increased by implementing a scheme in which antigens are multiplexed combinatorially^10^ (Klinger et al., 2015). Another difference is that, unlike antigen-binding or antigen-induced marker upregulation, the use of antigen-driven *in vitro* expansion is expected to select against anergic or regulatory cells that do not divide substantially upon stimulation, and instead highlight the most proliferative elements of the response. Clonal expansion also serves as a form of signal amplification to increase the sensitivity for rare clonotypes: a prior study reported a more sensitive detection of antigen-specific clonotypes when cells were isolated according to their antigen-driven proliferation (by dilution of a CFSE marker), compared to upregulation of an activation marker or binding to a peptide:MHC probe^25^ (Klinger et al., 2013).

Our CD40-targeted results indicate that stimulation with more immunogenic formulations of antigen can further increase the sensitivity with which rare clonotypes are detected. We show that anti-CD40 11B6-CD40L:Flu M1 immunogen elicits a response comprising substantially more detectable T cell clonotypes and homology clusters than the same protein in untargeted form, consistent with the observation that such targeting leads to increased T cell proliferation and cytokine production. This likely reflects a combination of CD40 activation of the APC concomitant with antigen uptake, focusing antigen to the APC via the anti-CD40 antibody binding, and specialized internalization into a dominantly early endosome compartment, resulting in sustained antigen presentation^21,26^ (Yin et al., 2016; Ceglia et al., 2021). As well as increasing the power to detect antigen-responsive TCRs, this likely provides a better (e.g., as compared to stimulation with peptide pools) representation of the response that is generated *in vivo* during natural infection or vaccination.

A limitation of the method we describe here is that it is unable to assign antigen specificity to antigen-expandable T cells that do not form receptor sequence homology clusters. Studies in which individual peptide:MHC-binding T cells were isolated and sequenced have defined, for most epitopes, a core group of TCRs belonging to one or more homology groups(s), and a remainder of TCRs that do not share evident sequence similarity^22,23^ (Dash et al., 2017; Glanville et al., 2017). This is consistent with a model in which a given 3-dimensional peptide:HLA antigen can often be recognized by a range of TCR sequence ‘solutions’ that are non-homologous in linear sequence space, but each of which can also tolerate some degree of homologous sequence variation. The magnitude of the T cell response corresponding to any given group is likely to be a function of both (i) the degree of sequence variation that is tolerated by the structure of the antigen, and (ii) the generation/maturation probability of TCRs within the group. The same considerations suggest that the homology groups to which our method is most sensitive (namely: groups that are frequently-generated and for which antigen-binding tolerates considerable sequence variation) are also the most likely groups to respond publicly across donors. Accordingly, we expect that future application of our method to larger cohorts will reveal that many of the identifiable CETs recur across individuals with matched HLA types.

The confident assignment of single chain TCR sequences to cognate antigens is complicated by several factors, including the heterodimeric nature of the TCR, the potential of any given TCR to cross-react with diverse antigens^27^ (Sewell, 2012), and the vast complexity of the repertoire found in any individual^28^ (Warren et al., 2011). Nonetheless, we show that unpaired α and β chains can be confidently assigned to antigen without cell isolation, and instead using statistical analysis of clonotype frequencies in expansion cultures. Moreover, our interrogation of unexpanded samples indicates that unpaired β chain clonotypes are sufficiently-specific biomarkers to enable inference of antigen-specificity within deep personalized repertoires down to frequencies less than 1e-6. The methodology developed here may be used to derive convenient high-fidelity biomarkers of antigen-specific T cell responses in the context of infection and/or vaccination studies. For example, applying this approach to longitudinal blood draws could enable highly-sensitive, multiplexed and antigen-resolved monitoring of the evolution of the circulating T cell response to a vaccine.

## Methods

### Anti-hCD40 monoclonal antibodies

Generation and screening strategies for making in-house recombinant anti-human CD40 12E12, 11B6 and 11B6-CD40L human IgG4 antibodies fused to dockerin at the H chain C-termini were as described^19,20,26^ (Flamar et al., 2012, 2013; Ceglia et al., 2021). Methods for expression vectors and protein production via transient or stable CHO-S (Chinese Hamster Ovary cells) transfection and quality assurance including CD40 binding specificity were as described^19,20,29^ (Flamar et al., 2012, 2013; Zurawski et al., 2017). Cohesin-Influenza Matrix 1 (Flu M1) protein has been described^19^ (Flamar et al., 2012).

### T cell expansion assay

Cryopreserved human PBMC from normal donors (AllCells, CA. ND1004 ID: 10504, ND1005 ID: 10002) were thawed with 50 U/ml benzonase^®^ nuclease (Millipore, cat 70746), washed and rested overnight in RPMI 1640 enriched with 1000 U/ml PenStrep (Gibco, 15140-122), 5 mM Hepes Buffer (Gibco, 15630-080), 1X Non-essential amino acids (NEAA) (Gibco, 11140-050), 1 mM Sodium Pyruvate (Gibco, 11360-070), 50 μM 2-Mercaptoethanol (Gibco, 21985-023), 2 mM Glutamax (Gibco, 35050-61) (herein called complete RPMI 1640) with 10% AB serum (GemCell, 100-512) in a 37°C 5% CO_2_ incubator. The following morning cells were cultured at a concentration of 2e6 cells/mL at 37°C in 1 mL complete RPMI 1640 + 10% AB serum in a 24 well flat bottom plate. Cells were treated with anti-CD40 non-covalently linked to a Cohesin Influenza Matrix1 (Coh-Flu M1) protein^19^ (Flamar et al., 2012), Coh-Flu M1 alone, or with 1 μM untargeted Flu M1 peptides (BEI Resources, Cat NR-21541), depending on the experiment. After forty-eight hours, 1 mL of complete RPMI 1640 with 10% AB serum and IL-2 (Proleukin, Sanofi) at a final concentration of 100 U/mL was added to each well. Half the media was changed at day 4 and at day 6 adding fresh IL-2. At day 10, cells were harvested and washed twice in PBS with 2 mM EDTA. For bulk RNA seq analyses, cells were spun down and the supernatant was removed to either store the cells at -80° C before proceeding with the analyses either as a pellet or to resuspend in RLT (Qiagen, cat 79216) + 1% 2-mercaptoethanol. For intracellular staining (ICS) or Luminex™ analyses, cells were instead resuspended in complete RPMI 1640 + 10% AB serum in 50 mL tubes, counted and rested over night at 37°C. The following day, cells were plated in a 96 well plate V bottom in 200 μL volume per well and re-stimulated with 2 μM Flu M1 peptides or controls for one hour in the case of ICS readout and up to 48 hours for Luminex™ analyses, at 37°C. Peptides were used in clusters named C1, C2 and C3 composed by, respectively, peptides 1-20, 21-40 and 41 to 60, or as single peptides or as small clusters of two or three overlapping peptides, used depending on the experiment. In the case of ICS, after one hour 0.175 μL of Golgi Stop (BD Golgi Stop, Cat 51-2092KZ) and 0.45 μL of Brefeldin A (BFA) (BD Cat 420601) were added and the cells were incubated for an additional 4 hours. Subsequently, cells were spun down and surface and intracellular staining was performed as described below gating on singlets, live cells, CD3^+^ followed by identification of TNFα^+^ and INFγ^+^ in both CD4^+^/CD8^-^ and CD4^-^/CD8^+^ cells. Cells analysed by Luminex™ were instead spun down after the re-stimulation time and the supernatant was analysed for secreted cytokines^30^ (Joo et al., 2014).

### Surface and Intracellular staining (ICS)

Human cells were first stained for surface markers. Human cells were transferred to a V bottom plate, washed twice in PBS and incubated for 20 minutes at 4°C with Live/Dead™ Fixable Aqua Dead Cell Stain Kit (Thermo Fisher Scientific, Cat. L34965) at a 1:50 dilution in a volume of 50 μL. Cells were washed twice with PBS and incubated for 30 minutes on ice with the mix of antibodies in a volume of 50 μL. After 30 minutes incubation on ice with the antibodies for surface staining, cells were washed in PBS twice and resuspended in Cytofix/Cytoperm™ (BD Biosciences) for 20 min at 4°C, followed by three washes in 1X Permwash (BD Biosciences). Cells were subsequently incubated at room temperature covered from light in 1X BD Permwash with the antibody mix for intracellular cytokines. Following the incubation time, cells were washed three times in 1X BD Permwash and resuspended in BD stabilizing fixative (BD Biosciences) diluted 1:3. All analysis plots were pre-gated on live (using Live/Dead stain) and singlet events. Cells were analyzed with a FACSCanto II or an LSR Fortessa (BD Biosciences). Data was analyzed with FlowJo® Software. The following antibodies were used: hCD3-BV711, clone UCHT1, ref 563725 (BD) or hCD3-PerCP clone SK7, ref 347344 (BD), hCD4-Pe-Cy7 clone SK3, ref 34879 (BD), hCD8-PacBlue clone 3B5, ref MHCD0828 (Invitrogen), hTNFα-APC clone RUO, ref 340534 (BD) and hINFγ-PE clone RUO, ref 340452 (BD).

### T cell receptor sequencing

RNA isolation from ND1001 (AllCells, CA, Donor ID: A5983), ND1002 (AllCells, CA, Donor ID: 9441), ND1004 and ND1005 cell pellets stored at -80° Celsius was performed using an AllPrep DNA/RNA Mini Kit (Qiagen). RNA quality was evaluated with an Agilent 2100 Bioanalyzer RNA pico kit (Agilent Technologies) prior to sequencing library preparation. T cell receptor sequencing libraries were prepared with the SMARTer Human TCR α/β Profiling Kit (catalog number 635015, Takara Bio USA, Inc.) according to manufacturer’s instructions with the exception of excluding the third and fourth bead size selection steps listed in Table 3 of the kit manual. Sequencing libraries were quantified using Kapa qPCR MasterMix (catalog number KK4973) on a QuantStudio7 Flex Real Time PCR System (Applied Biosystems by Thermo Fisher Scientific, Inc.). Libraries from different T cell cultures were pooled and 14 pM final library was added to the flow cell with 10% PhiX. Libraries were sequenced with MiSeq Reagent Kit v3 600 cycle (Illumina) to obtain 300 base-pair, paired-end reads.

### Analysis of TCR sequences

Raw sequencing data for each sample was mapped to germline segments using *mixcr* (MiLaboratory, version 3.0.11), to generate a clonotype list in which each entry is characterized by a unique combination of V and J segments and the CDR3 nucleotide sequence. For each sample, shortlists were constructed from the 1000 most frequent TCRα and TCRβ clonotypes, respectively, and all pairwise distance measurements were made on each shortlist using the TCRdist metric described previously^22^ (Dash et al., 2017). Hierarchical clustering was then performed on each set of distances using the *hclust* function in R, and clusters were identified at thresholds of 0, 5, 10, 15, 20, 25, 30, 35, 40, 45 and 50 using the *cutree* function. Each cluster was parameterized by its number of members, as well as the geometric mean frequency of its members. Significance was assigned to each cluster by determining the frequency with which clusters containing the same number of members, and a greater or equal mean frequency, were observed within 1000 random trials. Random trials used 1000 non-specific TCR clonotypes (of the corresponding chain), each assigned a frequency from the clonotype shortlist being tested, and clustered as described above. Clusters were considered significant at p<0.01 (i.e., <10 occurrences in the random trials). Clonotypes from significant clusters detected across all TCRdist thresholds were combined into a single master clonotype list and reclustered at the maximum threshold of 50 for final output. All code used for these analyses will be made available at https://github.com/TGenNorth/TCR_framework.git.

## Acknowledgments

We thank the Luminex core at Baylor Scott & White Research for performing the cytokine detection experiments and the Biotechnology core at Baylor Scott & White Research for developing the vaccine reagents. A Roche Collaborative Research Grant to the Baylor Scott & White Research Institute supported the initial development of anti-CD40 antibodies with enhanced agonist potency, and the Vaccine Research Institute via the ANR-10-LABX-77 grant funded the rest of the work.

## Authors contributions

**Valentina Ceglia**: Investigation, Methodology, Formal analysis, Data curation, Writing – original draft, review & editing. **Erin J Kelley:** Investigation, Methodology, Formal analysis, Data curation. **Annalee S Boyle:** Investigation, Methodology. **Yves Lévy**: Funding acquisition, Supervision, Visualization. **Gerard Zurawski**: Conceptualization, Methodology, Formal analysis, Data curation, Funding acquisition, Supervision, Writing - original draft, review and editing. **John A Altin**: Conceptualization, Methodology, Formal analysis, Data curation, Supervision, Writing - original draft, review and editing.

